# Quantitative assessment of angioplasty induced vascular inflammation with ^19^F cardiovascular magnetic resonance imaging

**DOI:** 10.1101/2022.11.25.518014

**Authors:** Fabian Nienhaus, Moritz Walz, Maik Rothe, Annika Jahn, Susanne Pfeiler, Lucas Busch, Manuel Stern, Christian Heiss, Lilian Vornholz, Sandra Cames, Mareike Cramer, Vera Schrauwen-Hinderling, Norbert Gerdes, Sebastian Temme, Michael Roden, Ulrich Flögel, Malte Kelm, Florian Bönner

## Abstract

Early macrophage rich vascular inflammation is a key feature in the pathophysiology of restenosis after angioplasty. ^19^F MRI with intravenously applied perfluorooctyl bromide-nanoemulsion (PFOB-NE) could offer ideal features for serial imaging of the inflammatory response after angioplasty. We aimed to non-invasively image monocyte/macrophage infiltration in response to angioplasty in pig carotid arteries using Fluorine-19 magnetic resonance imaging (^19^F MRI) to assess early inflammatory response to mechanical injury. Early macrophage rich vascular inflammation is a key feature in the pathophysiology of restenosis after angioplasty. ^19^F MRI with intravenously applied perfluorooctyl bromide-nanoemulsion (PFOB-NE) could offer ideal features for serial imaging of the inflammatory response after angioplasty. In eight minipigs, injury of the right carotid artery was induced by either balloon oversize angioplasty only (BA, n=4) or in combination with endothelial denudation (BA + ECDN, n=4). PFOB-NE was administered intravenously three days after injury followed by ^1^H and ^19^F MRI to assess vascular inflammatory burden at day six. Vascular response to mechanical injury was validated using immunohistology. Angioplasty was successfully induced in all eight pigs. Response to injury was characterized by positive remodeling with predominantly adventitial wall thickening and adventitial infiltration of monocytes/macrophages. ^19^F signal could be detected *in vivo* in four pigs following BA + ECDN with a robust signal-to-noise ratio (SNR) of 14.7 ± 4.8. *Ex vivo* analysis revealed a linear correlation of ^19^F SNR to local monocyte/macrophage cell density. Minimum detection limit of infiltrated monocytes/macrophages was as about 400 cells/mm^2^. Therefore, ^19^F MRI enables quantification of monocyte/macrophage infiltration after vascular injury with sufficient sensitivity. This might open an avenue to non-invasively monitor inflammatory response to mechanical injury after angioplasty and thus to identify individuals with distinct patterns of vascular inflammation promoting restenosis.

**One Sentence Summary:** ^19^F MRI enables radiation-free quantification of monocyte/macrophage infiltration after vascular injury with sufficient sensitivity.

## 1 Introduction

Despite advances in the development of drug eluting balloons (DEB) and stents (DES), restenosis after angioplasty remains a major challenge in daily clinical practice [1]. Proliferation of endothelial cells and smooth muscle cells lead to development of neointima and subsequent lumen narrowing. Emerging evidence suggest an important role of vascular inflammation early after angioplasty [2]. Infiltration of macrophages into the vascular wall precedes neointimal growth and quantity is directly correlated to neointima area [3]. Therefore, targeting macrophage polarization and infiltration is an unmet goal to prevent restenosis after angioplasty in peripheral artery disease (PAD) [4]. Imaging of monocyte/macrophage infiltration after angioplasty might identify patients at increased risk for restenosis early after endovascular intervention [5]. However, cell type specific positive contrast imaging of monocyte/macrophages and its association with restenosis after angioplasty using MRI has not been conducted so far.

We have shown, that ^19^F MRI is an emerging imaging technique that has proven its feasibility to visualize monocytes/macrophages in experimental disease models in mice at experimental field strength [6, 7]. This approach is based on the properties of monocytes/macrophages to ingest fluorine-containing perfluorocarbon-nanoemulsions (PFC-NE) which allows a specific cell tracking *in vivo* [8]. As the ^19^F signal is directly proportional to the amount of the PFC compound and there is a negligible natural ^19^F-background in the mammalian body, this imaging technique is highly specific and directly quantifiable by its positive contrast [6]. Recently, ^19^F MRI demonstrated feasibility using clinical scanners with sufficient signal-to-noise ratio (SNR) and reasonable scan time to reveal local inflammatory pattern after myocardial infarction [9–11]. Hence, ^19^F MRI may provide an ideal platform to visualize vascular inflammation and quantify monocytic cell accumulation *in vivo* after angioplasty of peripheral and central arteries. The goal of this study was to test SNR and minimal detection limits of ^19^F MRI with clinical applicable scanners at 3 Tesla to identify early monocyte rich inflammation in carotid arteries of pigs after graded vascular balloon injury.

## 2 Results

### 2.1 Experimental workflow and Safety of PFOB-NE application

Figure 1 shows the experimental workflow and protocol. Briefly, eight minipigs underwent graded carotid artery injury by either 15 min of oversized balloon angioplasty known to induce only subclinical stenosis [12] (BA, n=4) or a combination of 15 min oversized balloon angioplasty and mechanical denudation using a Fogarty catheter known to cause clinical significant stenosis [13] (BA + ECDN, n=4). All animals survived the procedure. Blood sampling and angiography was performed before and after vascular injury. At day three after injury PFOB-NE was administered intravenously. No side effects to PFOB-NE exposure could be recorded (for details see online data supplement). At day 6 after injury blood sampling as well as invasive angiography and IVUS was performed followed by *in vivo* and *ex vivo* ^1^H and ^19^F MRI and histology.

**Figure 1.**
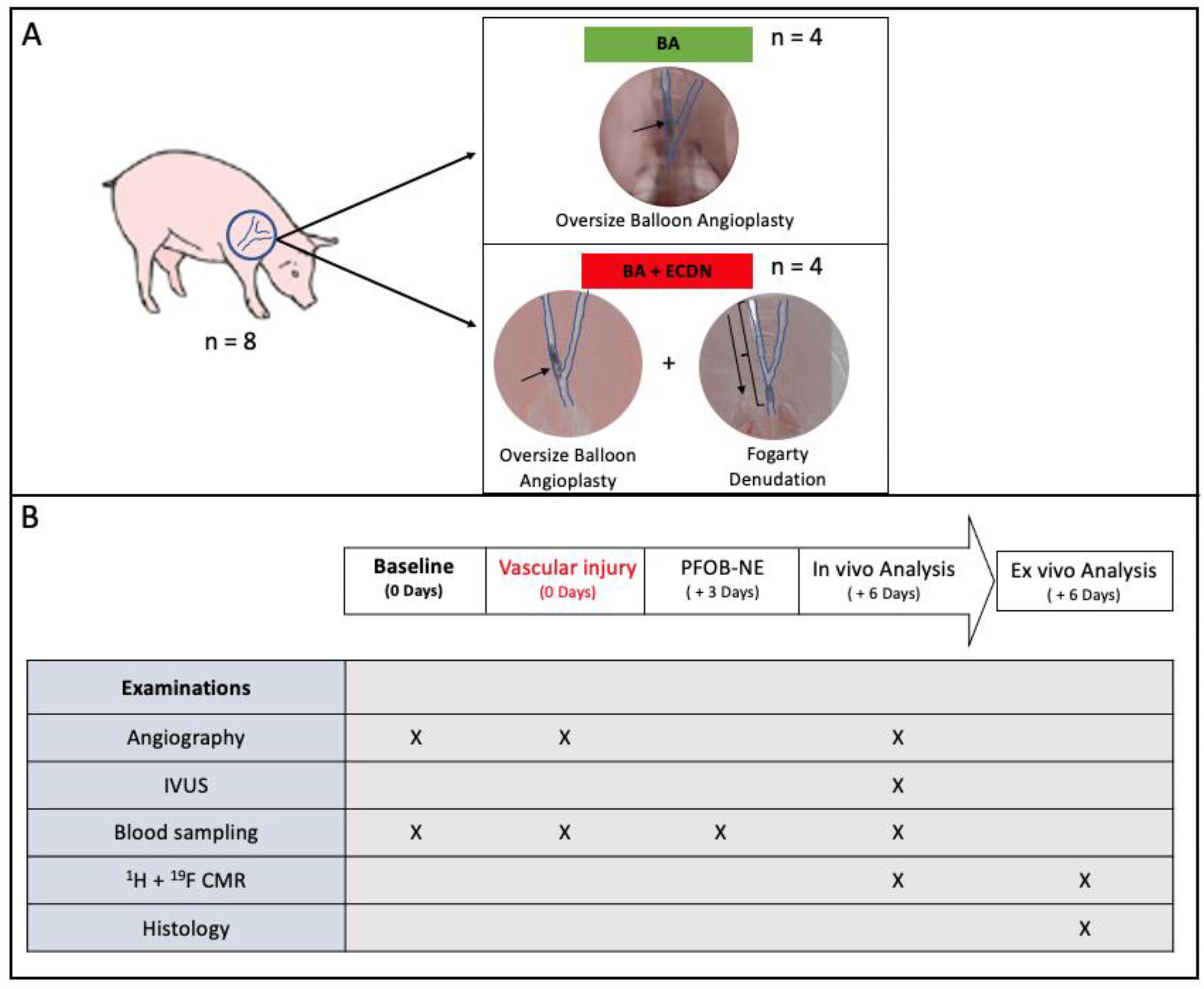
Experimental workflow and protocol. (**A**) Eight minipigs underwent either 15-min oversized balloon angioplasty (BA, n=4) or oversized balloon angioplasty with additional endothelial denudation (BA + ECDN, n=4) of the right carotid artery. (**B**) Vascular response to treatment was assessed by angiography at day zero. Perfluorooctyl bromide-nanoemulsion (PFOB-NE) was administered 3 days after injury. Vessel patency, analysis of the vascular wall and vessel blood flow was assessed by angiography and IVUS at day 6. Afterwards, pigs were examined for vascular inflammation using in vivo and ex vivo ^1^H/^19^F MRI followed by histology. *BA* = *Balloon Angioplasty; ECDN* = *Endothelial Denudation*

### 2.2 Effective induction of graded vascular injury and early remodeling

At day 7 after injury, angiographic assessment and IVUS revealed no relevant stenosis after BA. However, there was a 56% and 33% diameter stenosis with maintained blood flow in two pigs after BA + ECDN while two pigs suffered from nearly total carotid artery occlusion (**Fig. 2A**).

**Figure 2.**
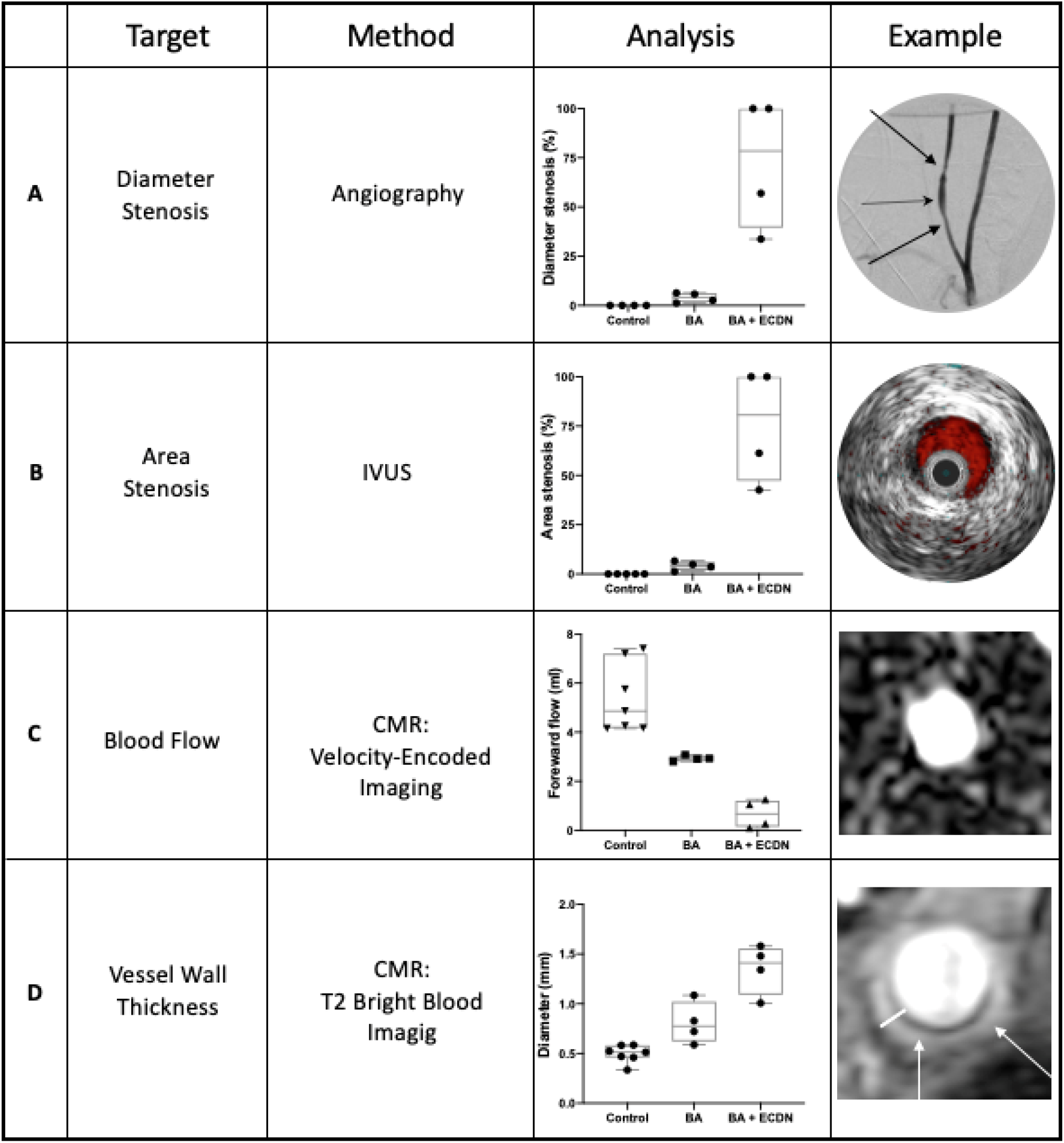
Angioplasty induced morphological and functional vascular adaptations. Carotid artery stenosis was evaluated using invasive angiography (**A**) and IVUS (**B**) Velocity-encoded MRI was used to quantify carotid blood flow (**C**) and vessel wall thickness was measured in MR-Angiography (**D**). Examples of the resulting images after BA + ECDN are shown on the right. BA + ECDN induced a relevant carotid artery stenosis or occlusion accompanied by reduction in blood flow and increased vessel wall thickness. *BA* = *Balloon Angioplasty*; *ECDN* = *Endothelial Denudation*

Carotid artery blood flow was assessed non-invasively by VENC MRI (**Fig. 2B**). Following BA + ECDN, a decrease in forward flow and post stenotic maximum velocity could be detected in all four pigs. Again, two pigs showed signs of nearly total artery occlusion with a decrease in forward flow and maximal velocity to nearly zero.

Vessel wall thickness was quantified non-invasively using high-resolution T2-weighted bright blood imaging (**Fig. 2C**). After BA, vessel wall thickness was found to be approximately 0.8 mm, versus 1.4 mm at day 7 after BA + ECDN.

### 2.3 Positive Vascular early remodeling after vessel injury

Histological hematoxylin and eosin (HE) staining was performed to quantify early vascular remodeling after graded vessel injury. Typical examples of the resulting images in 20x and 200x magnification are shown in **Fig. 3A**.

**Figure 3.**
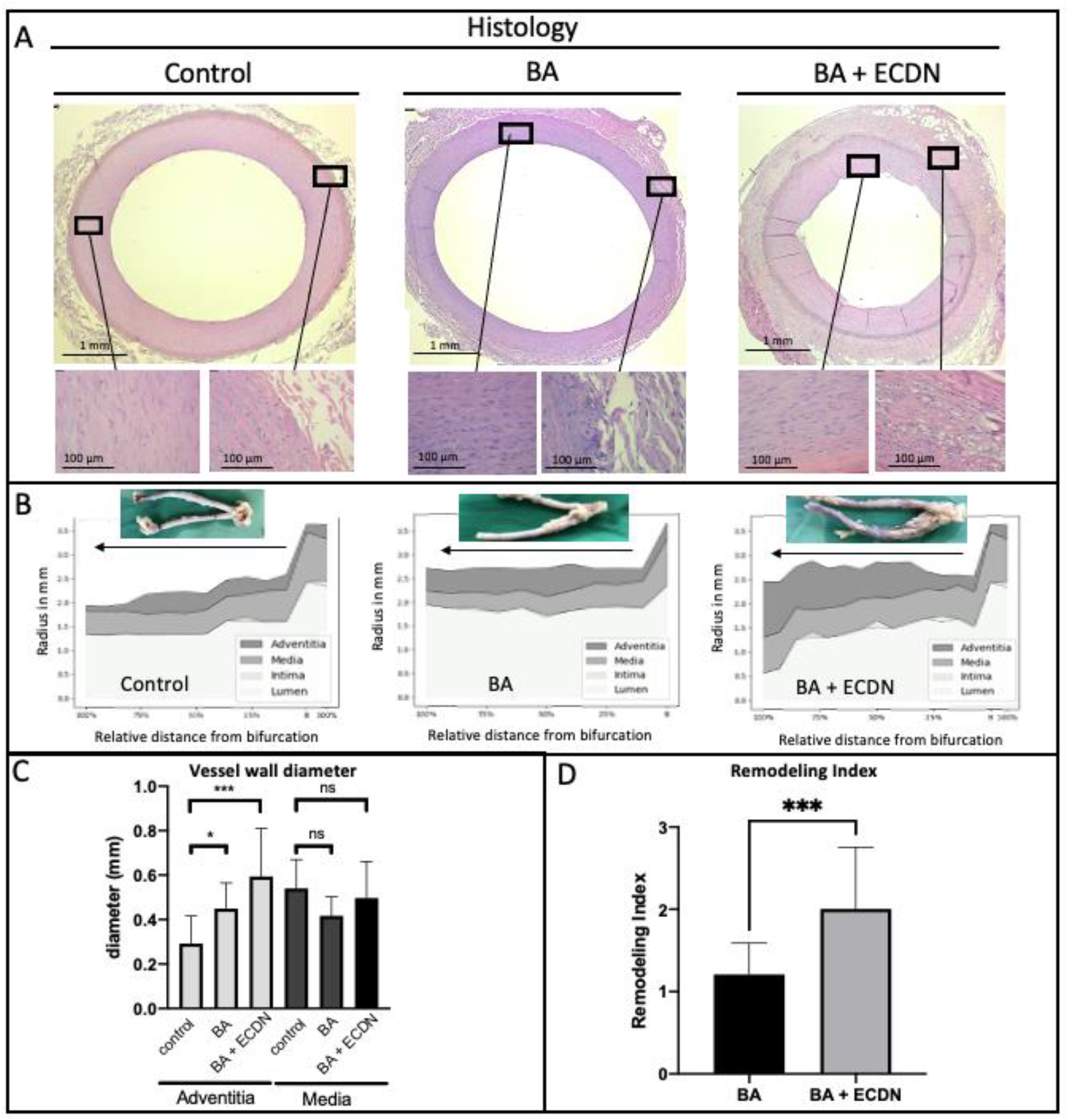
Histological analysis of carotid artery after angioplasty. Explanted carotid arteries of all 8 animals were cut into 15-20 sections. Analysis was performed in every resulting histologic sections and displayed as mean ± SD. (**A**) Typical examples of HE stained histological sections six days after graded vascular injury. Macroscopic images of the analyzed vessel are shown above. (**B**) Vascular lumen, intima, media and adventitia thickness along the entire vessel length as given in relative distance from bifurcation. (**C**) Measurements of vascular wall thickness of media and adventitia in BA and BA + ECDN as well as contralateral control carotid artery. (**D**) Remodeling index (External elastic lamina Area diseased/control) reveals positive vascular remodeling after both BA and BA + ECDN with aggravated remodeling after BA + ECDN. *BA* = *Balloon Angioplasty; ECDN* = *Endothelial Denudation; ns* = *not significant, *p<0.05, **p<0.01, ***p<0.001*,

Mean diameter of lumen, intima, media and adventitia was measured in at least 15 cross-sections starting from the carotid artery bifurcation and expressed as relative distance from the bifurcation (**Fig. 3B**). An increase in adventitial diameter compared to contralateral control carotid arteries could be documented after BA and BA + ECDN while diameter of media and intima remained unaffected. Only after BA + ECDN a relevant reduction of luminal radius was common. Quantification of vessel wall diameter across all histological sections obtained from all eight included pigs verified the findings in the individual arteries shown above (**Fig. 3C**). Compared to contralateral control carotid arteries, adventitial wall thickness increased 1,5-fold after BA and 2-fold after BA + ECDN, while media thickness was not significantly changed. Remodeling index (external elastic lamina area diseased/control) revealed significantly aggravated positive early vascular remodeling after BA + ECDN compared to BA (**Fig. 3D**).

### 2.4 ^19^F MRI reveals focal spots of monocyte invasion after injury

To assess inflammatory processes in addition to anatomical ^1^H MR images, ^19^F images were displayed on a ‘hot iron’ color scale and fused with ^1^H series to ensure exact anatomical localization of ^19^F signal. Typical examples of the resulting images *in vivo* and *ex vivo* are shown in **Fig. 4**.

**Figure 4.**
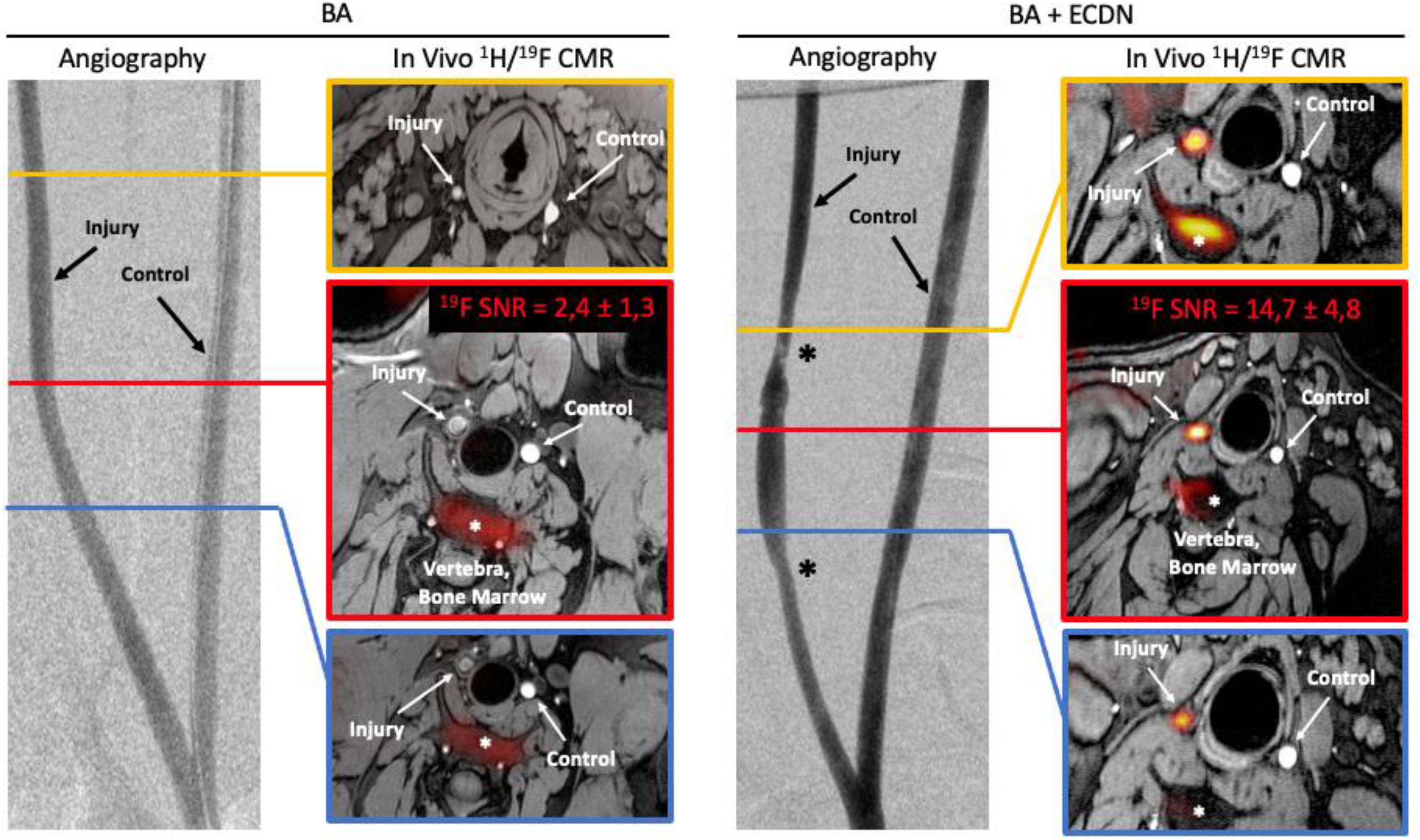
BA + ECDN induces a robust ^19^F signal in vivo. All pigs (n=8) were assessed for vascular ^19^F signal 6 days after graded vessel injury. Shown are angiographic images with indicated angioplasty regions of interest. Note a vasospasm (*) in the images of BA + ECDN. For anatomical *in vivo* co-localization ^19^F images were fused with a ^1^H 3D-T1 weighted images ^19^F signal intensity is displayed on a ‘hot iron’ color scale. After BA + ECDN, a robust ^19^F signal could be detected in all pigs (n=4) *in vivo*. No *in vivo* signal could be found in pigs following BA alone (n=4). *BA* = *Balloon Angioplasty; ECDN* = *Endothelial Denudation*

Following BA + ECDN, a robust *in vivo* ^19^F signal with a SNR of 14,7 ± 4,8 was detected in the treated carotid artery of all 4 pigs (**Fig. 4**). No signal was found in the contralateral control carotid artery. Vascular ^19^F signal could clearly be distinguished from background signals, mainly derived from the bone marrow and subcutaneous fat. No vascular ^19^F signal was found *in vivo* after BA. In the *ex vivo* setting at 3 T, ^19^F signal could be detected after 20 minutes in all 8 pigs with a strong SNR of 23,36 ± 3,8 after BA + ECDN and weak SNR of 6,2 ± 1,2 following BA. *In vivo* and *ex vivo* imaging revealed a patchy distribution of ^19^F signal along the carotid artery.

### 2.5 Local ^19^F signal co-registers with accumulation of monocytes/macrophages in a quantitative manner

To validate the patchy appearance of ^19^F signals and find minimal ^19^F detection limits for monocytes/macrophages, we conducted high resolution *ex vivo* ^1^H and ^19^F imaging at 9,4 T and 3 T and histological monocytes/macrophages staining (**Fig. 5A**). Fusion of ^1^H/^19^F images clearly demonstrates that ^19^F signal is derived from the vascular wall and that the ^19^F signal appears in a patchy fashion. Immunofluorescence staining confirmed a corresponding adventitial presence of monocytes/macrophages with predominantly CD163^-^CD68^+^ and to a lesser extend CD163^+^CD68^-^ appearance.

**Figure 5.**
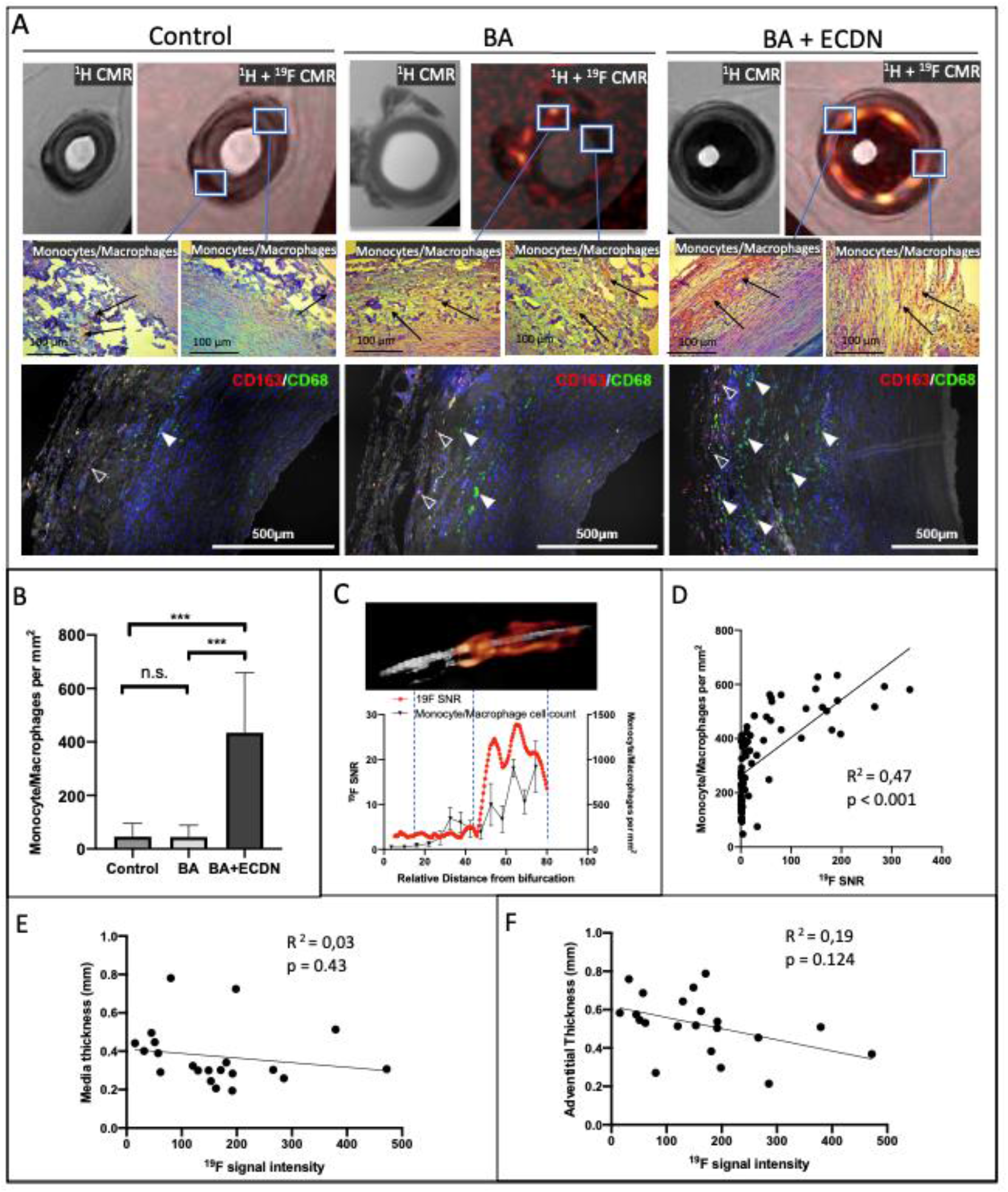
Local ^19^F signal co-localizes with accumulation of monocytes/macrophages independent of vessel wall thickness. High-resolution ex vivo ^1^H and fused ^1^H/^19^F MR images at 9.4 T show a robust ^19^F signal derived from the vascular wall after injury. Immunohistochemical staining for monocytes/macrophages was performed to characterize vascular immune cell infiltration 6 days after injury. Representative microscopy images from indicated MR regions. Immunofluorescence staining using CD163 (red, empty arrowheads) and CD68 (green, filled arrowheads) antibody confirmed a predominantly adventitial localization of monocytes/macrophages. (**A**). Monocytes/macrophages were counted from 15-20 histological cross sections per artery showing a large increase after BA + ECDN while the increase after BA remained only minor (**B**). Local ^19^F SNR measurements at 3 T were performed cross-sectional in correspondence to histologic cell count revealing an exact co-localization along the whole length of the artery (**C**) and 19F signal intensity correlates to monocyte/macrophage count (**D**). Moreover, ^19^F SNR does not correlate to (**E**) adventitia (**F**) and media thickness. *BA* = *Balloon Angioplasty; ECDN* = *Endothelial Denudation*

Semi-automated counting of monocytes/macrophages revealed a 17-fold increase in the adventitia after BA + ECDN while only a minor increase could be detected after BA **(Fig 5B)**. In the media, monocytes/macrophages were sparsely detected after both BA and BA + ECDN, suggesting that monocyte/macrophage infiltration early after vascular injury mainly occurs via the adventitia. The adventitial monocytes/macrophages phenotype was a mixture out of CD68^+^ CD163^-^ and CD68^-^ CD163^+^ cells with predominance of CD68^+^ CD163^-^ in BA + ECDN.

For histological validation carotid arteries were cut into at least 15 segments. Co-registration of local *ex vivo* ^19^F signal at 3 T with segmental monocytes/macrophages cell count was performed over the entire length of the carotid arteries of every included pig. This analysis of individual arteries revealing a close relationship between local ^19^F signal intensity as detected by MRI and spots of monocyte/macrophage accumulation. An individual example after BA + ECDN is shown in **Fig. 5C**. A highly significant correlation of local ^19^F signal with monocyte/macrophage cell density was found analyzing every segment of the treated carotid arteries after BA as well as BA + ECDN. The detection limit was about 400 monocytes/macrophages per mm^2^. No correlation could be found in relation to media **(Fig 5E)** or adventitial **(Fig. 5F)** wall thickness.

## 3 Discussion

Our study shows that 1) imaging carotid artery inflammation after balloon injury with ^19^F MRI is feasible in a reasonable time using a clinical scanner and a human size model; 2) local ^19^F signal co-registers with accumulation of monocytes/macrophages in a quantitative manner and 3) ^19^F signal reflects predominately early adventitial infiltration of monocytes/macrophages with CD 163^-^ CD68^+^ appearance associated with vascular wall thickening.

### Feasibility of ^19^F MRI to detect vascular monocytes/macrophages

Experimental approaches in mice have proven feasibility of ^19^F MRI to quantitatively visualize macrophage accumulation in a wide range of experimental set-ups and disease models, including vascular inflammation, [6, 7, 14]. However, those experiments were conducted with Perfluorpolyether (PFPE), which exerts excellent imaging properties, but a long body half-life making it less attractive for a potential clinical translation. In contrast, some PFCs, like PFOB, have a shorter biological half-life and have already been tested in clinical trials using their oxygen transport capacities. These compounds might have greater potential for clinical translation [15]. As has been shown recently, ^19^F MRI using PFOB nanoemulsion is also feasible in pre-clinical models after myocardial infarction using clinical scanners [9–11] with good agreement between hot spots of ^19^F signal and macrophage accumulation [11].

In the present study we used PFOB for ^19^F MRI in an animal model with a body size and an inflammatory reaction similar to humans to study macrophage visualisation after angioplasty. The resulting ^19^F signal in carotid arteries after balloon injury was highly reproducible with a robust *in vivo* and *ex vivo* SNR three or five times above the common detection threshold (SNR = 5). Therefore, our study provides evidence that ^19^F MRI using PFOB-nanoemulsion could be a valuable tool to monitor vascular monocytes/macrophages accumulation after angioplasty. Promising advances in MRI-technology such as improved coil designs and MRI sequence improvements have the potential to enhance the sensitivity of this approach. The current timing of PFOB-NE administration and ^19^F MRI imaging was adopted from our protocol of acute myocardial infarction [10, 11]. Since the time course of cellular response to vascular injury might be different to myocardial infarction, finding the optimal time point for PFOB-NE administration and ^19^F MRI imaging might even increase sensitivity of the method.

### Cellular Source of the ^19^F signal

We used a model of oversized balloon angioplasty with or without additional Fogarty-induced mechanical denudation (BA vs. BA + ECDN) according to established protocols [13]. A reliable and reproducible ^19^F signal *in vivo* could only be detected after BA + ECDN. The minimal detection threshold for detection of macrophages was about 300 to 400 cells/mm2. Untargeted perfluorocarbon nanoemulsions are taken up by M1 and M2 tissue macrophages to an equal extent [16]. Given the fact, that M1 macrophages were present in larger numbers in our model on day 6 after angioplasty, the ^19^F-signal observed reflected most likely M1 macrophages. BA + ECDN was always paralleled by adventitial wall thickening and positive early remodeling. In contrast, BA induced only a sparse infiltration of monocytes/macrophages in the vascular wall on immunohistology. This was paralleled by a small increase in adventitial wall thickening and early vascular remodeling index. The signal of those cells was only measurable under optimal coil distance conditions on *ex vivo* ^19^F scans.

Other studies using balloon oversize injury in pigs mainly focused on intimal hyperplasia, neointima formation and lumen area after a follow up of 2-4 weeks [12, 13]. Nevertheless, these studies confirm subclinical early vascular remodeling after overstretch injury with minimal reduction in lumen area [12]. In contrast, balloon oversize injury followed by endothelial denudation caused significantly reduced lumen area and enhanced early vascular remodeling [13]. Therefore, mild monocyte/macrophage infiltration under the ^19^F detection limit might also be of minor significance for early vascular remodeling and subsequent development of re-stenosis.

### Using ^19^F MRI to identify individuals at risk

Monocytes/macrophages play an important role in the pathophysiology of restenosis after angioplasty [3–5, 17]. Currently, there is no consensus for the default endovascular therapy with all commercially available interventions. There are suboptimal long-term outcomes due to high rates of re-stenosis and a discussion about apparently higher rates of death in patients treated with DEB [18–20]. However, manipulation of the monocyte/macrophage response after angioplasty could beneficially modulate restenosis rates in limb revascularisation [4].

Due to the insufficient treatment strategies so far, there is the medical need to identify patients at risk of restenosis after angioplasty. A prospective observational cohort study could demonstrate that imaging of vessel wall glucose uptake (^18^F-FDG) or microcalcification (^18^F-NaF) could predict restenosis following limb angioplasty [5]. Thus, already bulk identification of vessel wall surrogates of inflammation predicts restenosis. We have shown, that ^19^F MRI with an untargeted nanoemulsion can directly image and quantify monocytes/macrophages. The specificity of this approach to target M1 and M2 macrophages can even be improved by functionalizing the nanoemulsion. As has been shown in mice, multitargeted nanoemulsions with different ^19^F-agents identified by multi chemical shift selective imaging were capable of simultaneously revealing a broad range of antigens important in the development and exacerbation of atherosclerosis (“multi-colour-imaging”) [21]. This imaging platform could provide a complete phenotyping of vascular healing after angioplasty with the major advantage of a positive, specific and quantifiable signal with a. In the present study, using an untargeted nanoemulsion, we were able to directly visualize the monocyte/macrophage lesion burden in a quantitative manner with a resolution sufficient for improved vascular-wall mapping of the signal.

Interestingly, the minimal detection limit of 19F MRI was comparable to cell density of human carotid atherosclerotic macrophage-rich vulnerable plaques [22, 23]. As has been shown for primary prevention, the indirect visualization of inflammatory foci by enhanced glucose uptake using ^18^F-fluorodeoxyglucose (^18^F-FDG) predicts subsequent vascular events in carotid artery atherosclerotic lesions [24, 25]. Inflammatory microcalcification as identified by ^18^F-sodium-fluoride (^18^F-NaF) has already shown predictive value for cardiovascular events in the coronary arteries [26, 27]. Given the fact, that the number of monocytes/macrophages is directly related to plaque vulnerability [22, 28], quantitative imaging of monocytes/macrophages may have additional value for identifying the patient at risk.

### Study limitations

Our study only evaluates acute vascular inflammation. No statement can be made about the significance of ^19^F signal intensity for long-term vascular remodeling or chronic inflammatory processes in atherosclerosis. Further long-term studies with graded vascular injury and large animal models of atherosclerosis are needed to investigate vascular remodeling and plaque progression in relation to ^19^F signal intensity.

### Conclusion

^19^F MRI enables quantification of monocyte/macrophage infiltration after vascular injury with sufficient sensitivity. This might open an avenue to non-invasively monitor inflammatory response to mechanical injury after angioplasty and thus to identify individuals with distinct patterns of vascular inflammation promoting restenosis.

## 4 Methods

### 4.1 Model of Vascular Injury

The experiments were performed in eight adult Aachen minipigs [29] with a medium age of 2 years (± 5 months) which were breed, housed and extensively characterized [11] at the central animal facility center of Heinrich-Heine-University, Düsseldorf, Germany. All study protocols were carried out in accordance to the national guidelines of animal care and approved by the state authority ‘Landesamt für Natur-, Umwelt-und Verbraucherschutz’ (84-02.04.2018.A154 und 81-02.04.2019.A379). Eight heparinized pigs underwent graded carotid artery injury. Four pigs were subjected to 15 min of oversized balloon angioplasty only (Passeo-35, BIOTRONIK, Berlin, Germany) with a balloon to artery ratio of 1.3 : 1 known to induce only subclinical vascular changes with an area stenosis of 10-20% after 4 weeks (BA, n=4) [12]. The other four pigs underwent a combination of oversized balloon angioplasty and endothelial denudation using a Fogarty catheter known to induce a significant further decrease in lumen area and increased plaque size (BA + ECDN, n=4) [13]. Models were modified according to previous protocols [12, 13] and described in detail in the data supplement.

### 4.2 Production and Application of PFOB-Nanoemulsion

Perfluorooctyl bromide-nanoemulsion (PFOB-NE) was produced according to established protocols [30]. PFOB-NE was administered intravenously body weight adjusted (5 ml/kg BW) at day 3 after vascular injury according to the maximum circulating monocyte count and protocols adopted from experiments in myocardial infarction [11]. A detailed description of production and administration can be found in data supplement.

### 4.3 Invasive Assessment of Vascular Morphology

Before and seven days after vascular injury pigs were subjected to invasive assessment of carotid arteries. A detailed experimental flow chart and protocol is given in Figure 1 and data supplement. Briefly, carotid artery blood flow and vessel diameter were evaluated at baseline and six days after vascular injury by invasive angiography and intravascular ultrasound (IVUS). Serial blood samplings at day one, three and seven was conducted to investigate circulating inflammatory response to injury.

### 4.4 ^1^H and ^19^F MRI data acquisition

Immediately after vascular assessment at day 6, non-invasive ^1^H and ^19^F MRI was performed using a whole-body 3.0 T Achieva X-series scanner (Philips Healthcare, Best, the Netherlands). The MRI scan consisted of a ^1^H protocol including 3D time of flight (TOF) angiography, phase-contrast velocity encoded (VENC) measurements and high-resolution T2-weighted black blood sequences for vessel wall assessment. This was followed by a ^19^F protocol for monocyte/macrophage imaging with tested and optimized sequences [10]. After acquisition of ^1^H reference scans, the pigs were removed from the bore, and the ^19^F coil was placed on the neck directly above the injured artery. Pigs were repositioned for optimal dataset overlay as has been tested before for local signal precision [10].

### 4.5 MRI data analysis

MRI datasets were visualized and analyzed for signal overlays using HOROS (Nimble Co LLC, USA) and Circle CVI (Circle Cardiovascular Imaging Inc., Calgary, AB, Canada) for analysis of anatomic dimensions and velocities.

For assessment of ^19^F datasets, exact anatomical colocalization was performed by image fusion of ^1^H and ^19^F datasets using HOROS. For quantification of ^19^F signal intensity, the primary signal at a respective region was corrected according to the coil sensitivity profile as published previously [11]. Then, SNRs were calculated from the ratio of the mean of a region of interest (ROI) and the standard deviation of the noise of a ROI outside the body [10, 11]. After in vivo ^1^H and ^19^F MRI acquisitions, pigs were euthanized with potassium chloride and Narcoren (Pentobarbital sodium®, Boehringer Ingelheim Vetmedica, Ingelheim, Germany) inside the scanner.

### 4.6 Autopsy, ex vivo MRI and histology

Autopsy was performed and both carotid arteries and the carotid trunk were excised *in toto* and stored in 4%paraformaldehyde (PFA) for at least 7 days. Paraffin embedded vessels were sectioned (5 μm) and stained with Hematoxylin/Eosin (H/E). For immunoflorescence and immunohistology the following antibodies were used: anti-CD163 antibody (clone 2A10/11, Bio-Rad Laboratories, Inc., Hercules, California, USA); anti-CD68 antibody (ab125212, 7.5μg/ml, abcam, Cambridge, UK). Tissue processing is described in detail in the data supplement. Histological examination was conducted with a Leica microscope (DM 4000 M, Leica Microsystems, Wetzlar, Germany) with 10×, 20× and for detailed images 40x magnifying lenses. H/E or CD163/CD68 stained cells were counted semi-automatically using ImageJ (LOCI, Wisconsin) with a detailed protocol provided in the data supplement. Early Remodeling Index (RI) was calculated as lesion site divided by untreated control vessel external elastic membrane (EEM).

### 4.7 Statistical Analysis

Statistical analysis was performed using GraphPad Prism (Version 9, IBM, San Diego, CA, USA). Unless otherwise stated, continuous variables are presented as mean ± standard deviation (SD). Normal distribution was tested using the Shapiro-Wilk test. Data between the two different groups (BA and BA + ECDN) were analyzed by 2-sided unpaired Student’s t-tests for normally distributed data and Mann-Whitney U-test for not normally distributed data. Pearson’s or Spearman’s correlation was used to assess the relationship between different MRI parameters.

## List of Supplementary Materials

Materials and Methods

## Acknowledgments

We thank Juliane Geisler, Christoph Jacoby and Martin Sager for their excellent technical and administrative support.

## Funding

This study was supported by the German Research Foundation SFB 1116 projects B12 (M.R., M.K.), B10 (U.F.), Gerok Scholarship (F.N.), TRR 259 project B03 (F.B., U.F.) and project grants BO4264/1-1 (F.B.), TE1209/1-1 (S.T.), FL303/6-1 (U.F.) as well as by grants from the German Federal Ministry of Health (BMG) and Ministry of Culture and Science of the State North Rhine-Westfalia (MKW NRW) to the German Diabetes Center (DDZ) and by the Federal Ministry of Education and Research (BMBF) to German Center for Diabetes Research (DZD e.V.) (all M.Rot., S.C., V.S.-H., M.Rod.) and the Schmutzler Stiftung (M.Rod.).

## Author contributions

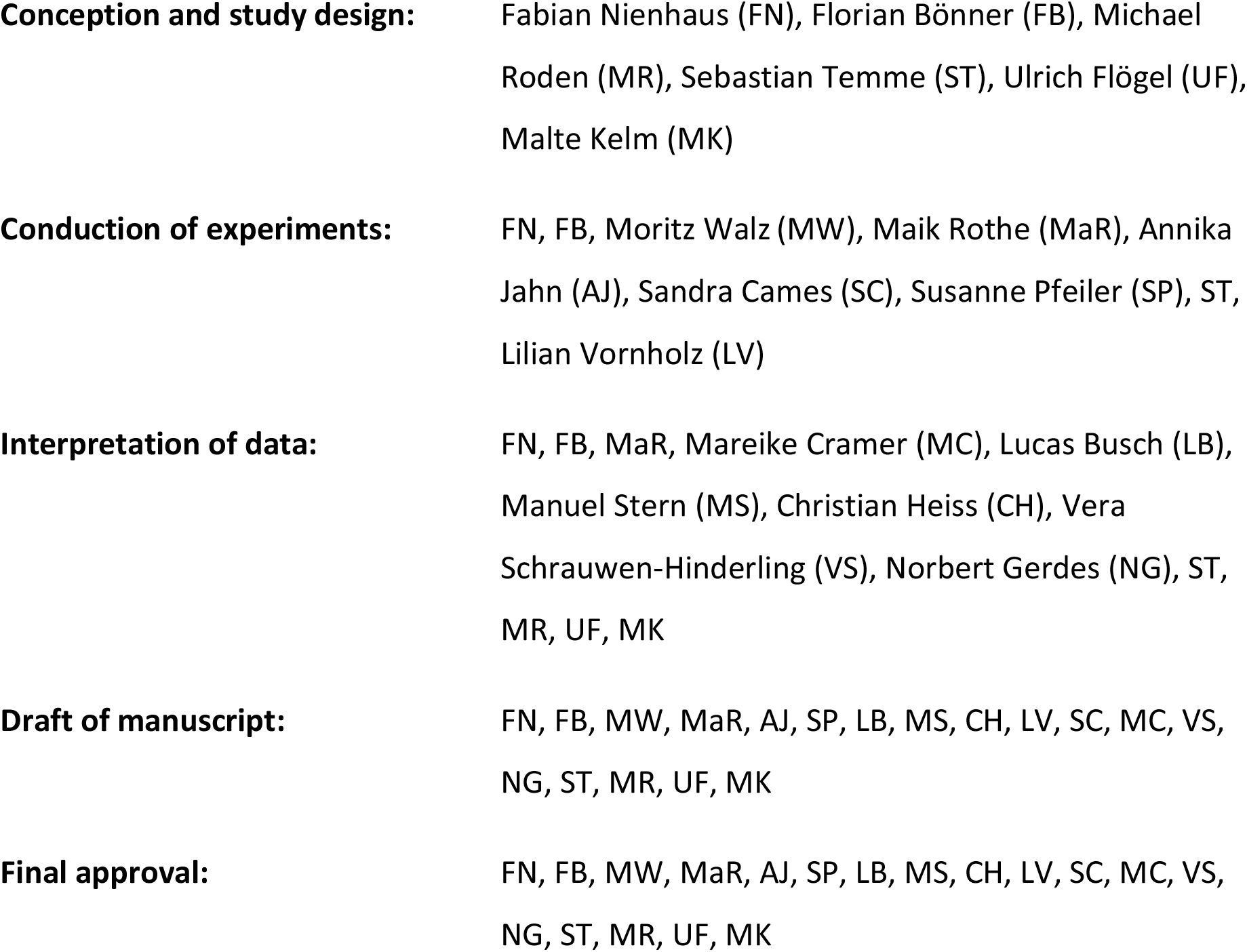

## Competing interests

None

## Data and materials availability

The data that support the findings of this study are available from the corresponding author, FB, upon reasonable request.

## Notes

### Competing Interest Statement

The authors have declared no competing interest.

